# Caramel Dye IV Induces Oxidative Stress Damage In Liver And Kidney From Mice

**DOI:** 10.1101/824276

**Authors:** Emerson Marins, Julia Spanhol da Silva, Pâmela Carvalho da Rosa, Vitor Antunes de Oliveira, Aline Zuanazzi Pasinato, Joana Grandó Moretto, José Eduardo Vargas, Felix Alexandre Antunes Soares, Rômulo Pillon Barcelos

## Abstract

Caramel dye IV (C-IV) is a synthetic organic product, does not present nutritional, ergogenic, or technological factors, but leads to reactive oxygen species (ROS), causing damage to a wide range of molecules, leading to cancer, cardiovascular and neurodegenerative diseases development. We aimed to verify the effects of different doses of C-IV dye on the markers of oxidative stress in the liver and kidneys from male Swiss CF-1 mice, divided into 4 experimental groups: control; C-IV 0.3g/kg; C-IV 1g/kg and C-IV 3g/kg. We found that 3 g/Kg of C-IV dye promote oxidative damage in liver and kidney homogenates, evidenced by the increase of lipid peroxidation, reduction of free SH groups, and higher ROS production. As a consequence, increased superoxide dismutase and acetylcholinesterase enzymes activities were detected. These damages were confirmed through histology images. These results indicate that daily doses might induce oxidative stress damages and possible lead to chronic diseases development.

## 1. INTRODUCTION

Dyes are usual ingredients used in food products and generally do not have nutritional, ergogenic, or technological importance for consumers, is used for food aesthetic purpose ^1^. Among the dyes allowed as a food additive, (C-IV) has a special place among dyes as being the oldest and the most commonly used by the food industry in a wide range of foods ^2^, and is classified as a synthetic organic product (ANS), 2010).

This additive is produced through sulphite-ammonia process; some acids and alkalis may also be added during its production ^4^, as sulfuric or citric acid, phosphoric, acetic and / or carbonic acid; alkalines such as hydroxides or mixtures of sodium, potassium and / or calcium; and salts such as carbonate, hydrogen carbonate, sulfate and ammonium, phosphate sodium, potassium or calcium. Some ammonium compounds, such as carbonate and ammonium hydrogen carbonate may also be added ^5^. In this process, an ammonia interaction leads to sugar reduction, and byproducts such as 4-methyl imidazole (4-MEI) generation, which might presents highly toxic effects, such as neurological and cancerous pathologies ^6^ which lead, for example, the State of California to classify ut as a “cancer-causing chemical”^5^. According to the Expert Committee on Food Additives (JECFA) determined an Acceptable Daily Intake (IDA) of 300mg / kg p.c. of Caramel IV dye ^5^.

A toxic intake can develop signs and symptoms that reveal an imbalance produced by the interaction of the toxic agent with the organism ^7^. Such imbalance can occur due to free radicals overproduction, that leads a pathological process named oxidative stress if that exceeds the antioxidant defenses ^8^. Oxidative stress causes damage to a wide range of molecules including lipids, proteins, and nucleic acids that might lead to the development of diseases such as cancer, cardiovascular and neurodegenerative diseases ^9^.

Therefore, an investigation of possible toxic effects of C-IV dye has great importance, since it is included in a lot of food and drinks and, this way, consumed daily worldwide, mainly due to the lack of knowledge of the exact mechanisms responsible for its harmful effects and toxicity. Thus, this work aims to verify the effects of different doses of Caramel IV dye on the markers of oxidative stress in the liver and kidney of mice.

## 2. METHODS

### 2.1 Animals

Thirty-six male Swiss CF-1 male mice (30-40g) g were obtained from our own breeding colony and kept in plastic boxes containing a maximum of five animals per cage under controlled environment conditions (12:12h light-dark cycle, with onset of light phase at 7:00, 25±1 °C, 55% relative humidity) with standard food and water ad libitum. Mice were acclimated for 7 days before initiation of any procedures. All experiments were conducted following national and international legislation: Brazilian College of Animal Experimentation (COBEA) and the U.S. Public Health Service’s Policy on Human Care and Use of Laboratory Animals-PHS Policy and with the approval of the local Ethics Committee (#013/2017).

The animals were randomly assigned into four groups and submitted to single and individual saline administration or Caramel IV dye by intra-gastric tube and kept in cages with only water.

### 2.2 Experimental protocol

Caramel IV (C-IV) was purchased from Prime Foods São Paulo, Brazil (lot 120117C), dissolved in saline and administered via intra-gastric gavage (i.g). Animals were kept in a 12 hours fast, randomly divided into four groups - nine animals per group: (A) control saline; (B) C-IV 0.3 g/kg; (c) C-IV 1g/kg; (d) C-IV 3 g/kg. These dosages were based on Recommended Dietary Allowance (RDA) (300mg/kg per day to humans) ^10^. Thus, the highest dosage used in this study is equivalent to 3,000mg/kg/day for ingestion of Caramel IV dye, 10 fold higher than the RDA. To study the effects of C-IV the i.g. route of administration was chosen since it is the most commonly used way that this dye is consumed worldwide.

After 4 hours of administration of the different doses, the animals were anesthetized with ketamine and xylazine for getting and separation of plasma through cardiac puncture and after euthanized by cervical dislocation. Liver and kidneys were removed and separated immediately. The experimental design is illustrated in Figure 1.

**Fig. 1.**
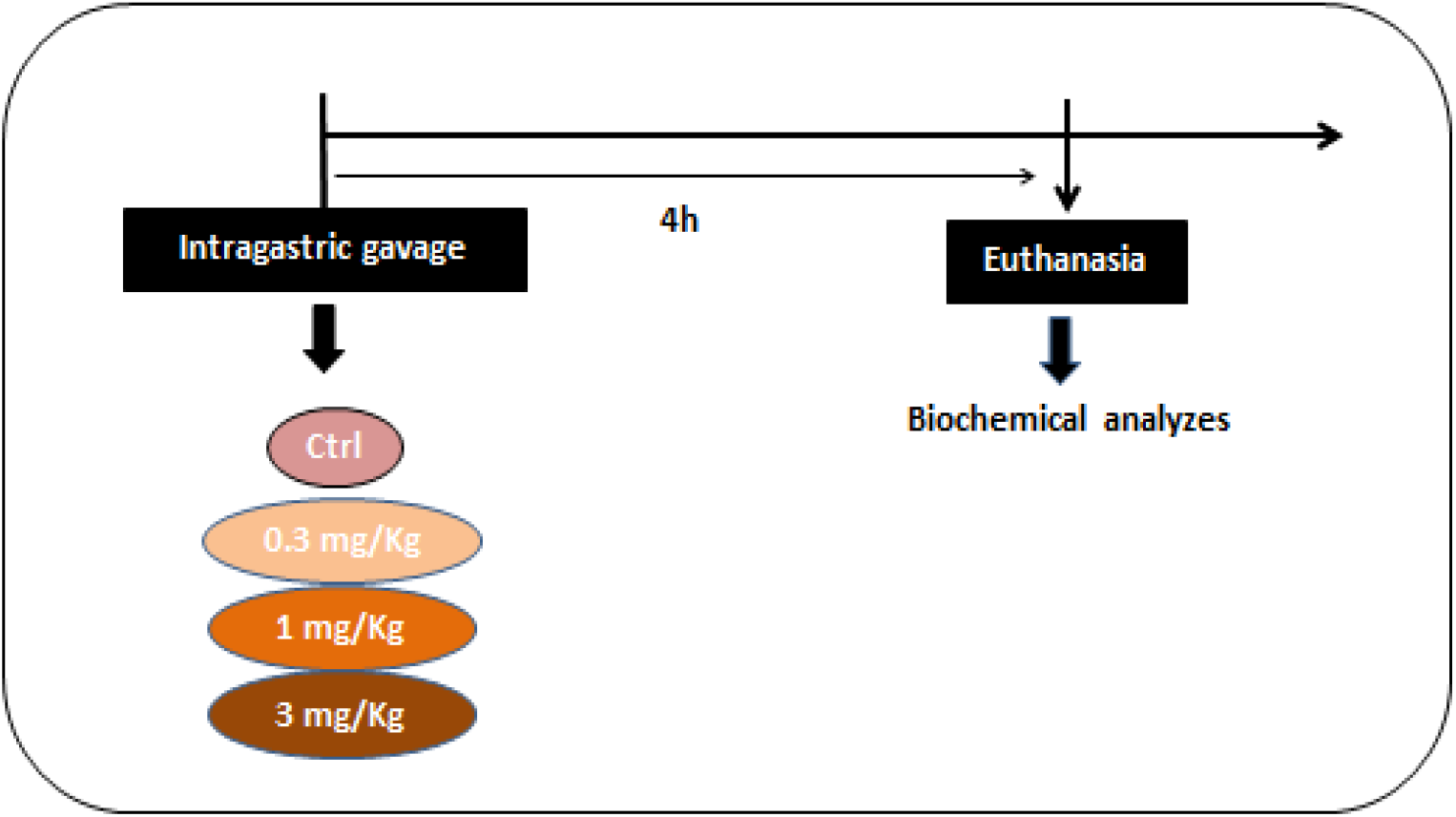
Description of experimental protocol used in this study.

### 2.3 Oxidative Stress Markers

#### 2.3.1 Lipid Peroxidation assay

Lipid peroxidation was estimated by quantifying thiobarbituric acid reactive substances (TBARS). Aliquots (200μL) of liver and kidney homogenates were mixed with 500μL thiobarbituric acid (0.6%), 200 μL sodium dodecyl sulfate (SDS) 8.1%, and 500 μL acetic acid (pH 3.4) and incubated at 90° C for 1 h. TBARS levels were measured at 532 nm (Spectrophotometer Proanalysis, UV / VIS 190-1100nm UV-1600) using a standard curve of malondialdehyde (MDA), and the results were reported as % of control ^11^.

#### 2.3.2 ROS production assay

ROS generation was determined spectrofluorimetrically in liver homogenate using 2′7′dichlorodihydrofluorescein diacetate (H_2_DCF-DA) (1 mM). The oxidation of H_2_DCF to 2′7′dichlorofluorescein (DCF) is used as an index of the peroxide production by cellular components ^12^. Briefly, liver and kidney homogenates were added to the standard medium, and the fluorescence was determined at 488 nm for excitation and 525 nm for emission, with slit widths of 3 nm. Oxidized dichlorofluorescein was determined using a standard curve of oxidized dichlorofluorescein, and the results were expressed as oxidized DCF μmol/mg protein ^13^.

#### 2.3.3 Non-protein thiol measurement (NPSH)

To estimate GSH content, we determined NPSH as follows: 100 μL of 10% TCA was added to 100 μL of either the S1 homogenates of liver or kidney. After centrifugation (4,000 × g at 4°C for 10 min), the protein pellet was discarded and free –SH were determined in the clear supernatant (which was previously neutralized with 0.1 M NaOH) according to Ellman (1959) (412 nm – Spectrophotometer Proanalysis, UV / VIS 190-1100nm UV-1600)

### 2.4 Antioxidant defense enzymes

#### 2.4.1 Catalase Activity (CAT)

The activity of the CAT enzyme was determined according to the method proposed by Aebi (1984). Briefly, S1 aliquot (50 μL) was added to a medium containing potassium phosphate buffer (50 mM; pH 7.4) and H_2_O_2_ (1 mM). The kinetic analysis of CAT was started after H_2_O_2_ addition, and the color reaction was measured at 240 nm. One unit of the enzyme is considered as the amount which decomposes 1 μmol H_2_O_2_/min at pH 7

#### 2.4.2 Superoxide desmutase activity (SOD)

The activity of the SOD enzyme was determined according to the method proposed by Misra and Fridovich (1972). The kinetic analysis of SOD was initiated after the addition of adrenaline, and the color reaction measured in a spectrophotometer at 480 nm. Data were corrected for protein content determined, according to Bradford (1976) using albumin as standard, and expressed as U SOD/mg protein ^17^.

#### 2.4.3 Acetylcholinesterase

The AChE activity was estimated in liver, kidney or plasma by the Ellman method ^18^ using acetylthiocholine iodide (ATC) as substrate and etopropazine as butyrylcholinesterase (BChE) inhibitor ^19^. Briefly, 880μl of 0.1M TFK buffer, added with 25 μl of liver or kidney homogenized tissue, was used and exposed to the water bath for 2 minutes at 30° C. Then, 50 μl of ATC 9 mM and 50 μl of DTNB 6 mM were added for readings at 412 nm at the 0, 60 and 120 second times, being represented by nmol of ATC min/mg.

#### 2.4.4. Kidney and Liver Histopathological Analyses

After euthanasia liver and one kidney were rapidly extracted and fixed in 4% paraformaldehyde for 24 hours and then paraffin embedded for histopathological and analyzed ^20^. The samples were embedded in paraffin and sectioned using a microtome (5 mm of section thick). For histopathological evaluation, alternate sections were deparaffined, rehydrated, and finally stained with hematoxylin-eosin according to standard procedures^21^. The sections were then observed with light microscopy at a final magnification of 400x (Olympus, Germany).

### 2.5 Statistical analysis

Statistical analysis was performed using the GraphPad Prism 6^®^ program, with one-way analysis of variance (ANOVA), followed by Tukey post-test. Data are expressed as means ± Standard Error of Mean (S.E.M.), p values < 0.05 were considered significant.

## 3. RESULTS

### 3.1 Markers of oxidative damage

#### 3.1.1 Lipid peroxidation

Groups C-IV 1g/kg and C-IV 3g/kg increased TBARS levels in the liver when compared to the control and C-IV 0.3g/kg groups (Figure 2A, p<0.0027). No significant changes were observed in mice’s kidney (Figure 2B).

**Fig. 2.**
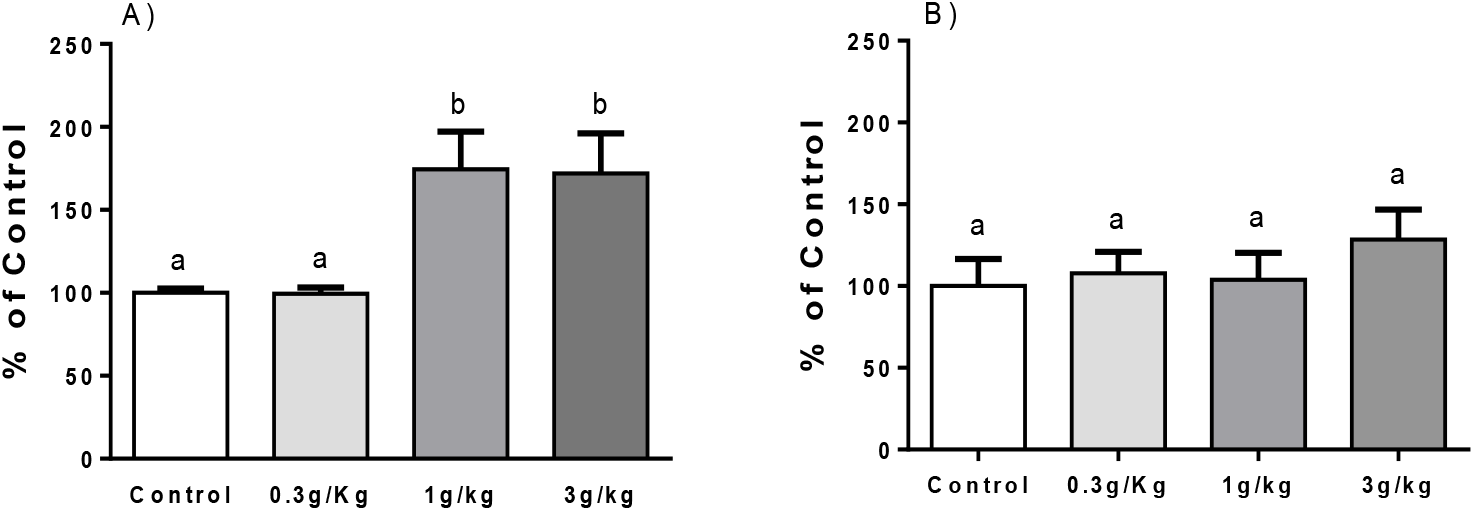
Effect of Caramel IV on lipid peroxidation (TBARS levels) in liver (A) and kidney (B) of mice. Data are represented as % of control and expressed as mean ± S.E.M (n = 9, p<0.05). Means for a variable with superscripts without a common letter differ.

#### 3.1.2 ROS production

No significant difference was observed on the liver (Figure 3A). However, an increased ROS production was observed at C-IV 3g/kg group when compared to all other groups in the kidney (Figure 3B, p<0.0049).

**Fig. 3.**
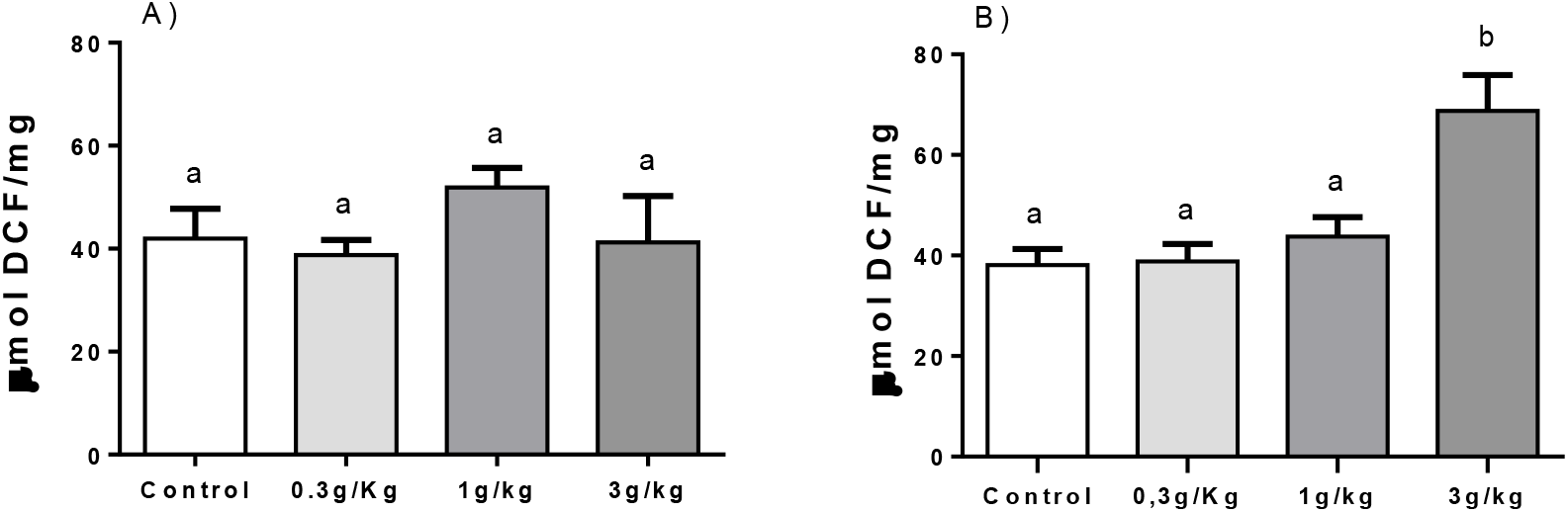
Effect of Caramel IV on ROS production in liver (A) and kidney (B) of mice. Data are represented as μmol of DCF/mg and expressed as mean ± S.E.M (n = 9, p<0.05). Means for a variable with superscripts without a common letter differ.

#### 3.1.3 Non-protein thiol measurement (NPSH)

No changes were observed between groups on the liver (Figure 4A). However, a reduce NPSH levels was observed in the C-IV 3g/kg group when compared to the C-IV 1g/kg group on the kidney of mice (Figure 4B, p<0.05).

**Fig. 4.**
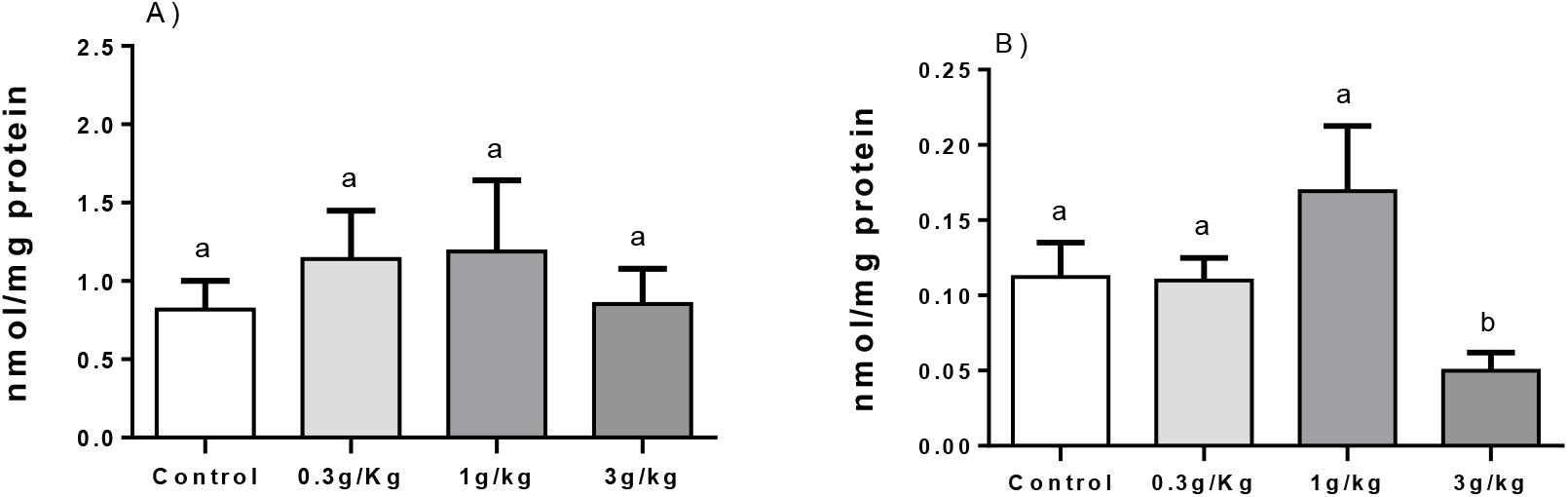
Effect of Caramel IV on NPSH levels in liver (A) and kidney (B) of mice. Data are represented as nmol/mg protein and expressed as mean ± S.E.M (n = 9, p<0.05). Means for a variable with superscripts without a common letter differ.

### 3.2 Antioxidant defense enzymes

#### 3.2.1 Catalase (CAT) activity

No significant difference was observed in the CAT activity in the hepatic tissue (Figure 5A). However, increased CAT activity was observed on the kidney at C-IV 3g/kg groups when compared to the other groups (Figure 5B, p<0.0267).

**Fig. 5.**
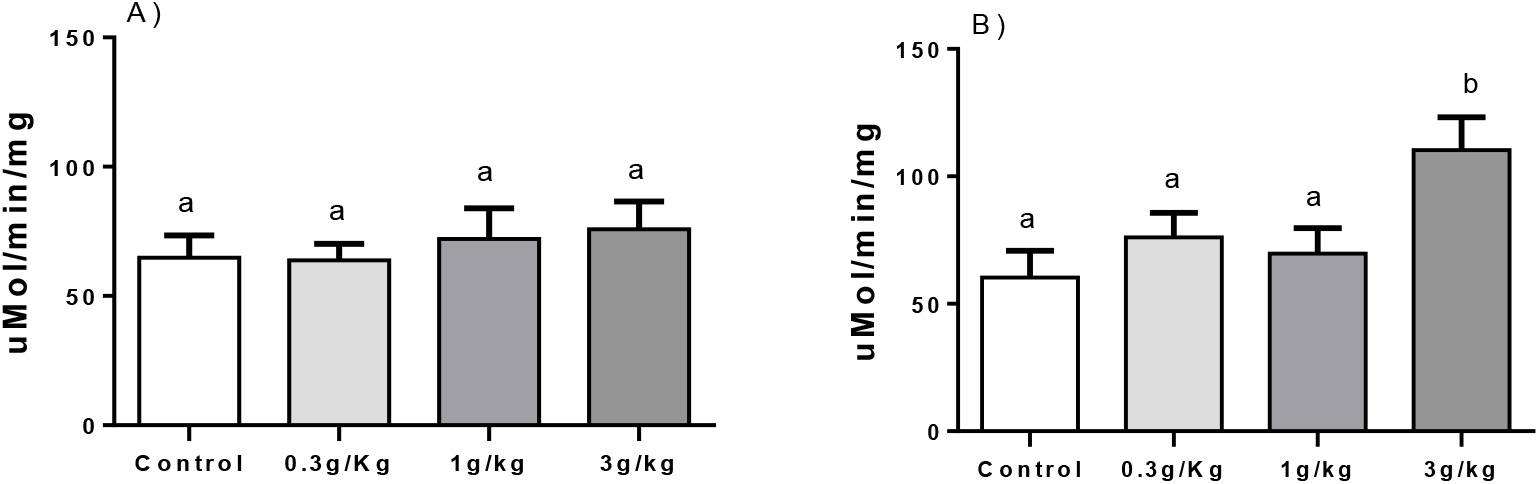
Effect of Caramel IV on catalase activity in liver (A) and kidney (B) of mice. Data are represented as μmol/min/mg and expressed as mean ± S.E.M (n =9, p<0.05). Means for a variable with superscripts without a common letter differ.

#### 3.2.2 Superoxide dismutase (SOD) activity

We observed an increased SOD activity in the C-IV 3g/kg group when compared to the other groups both on liver and kidney of mice (Figure 6A, p<0,0013 and 6B, p<0.05).

**Fig. 6.**
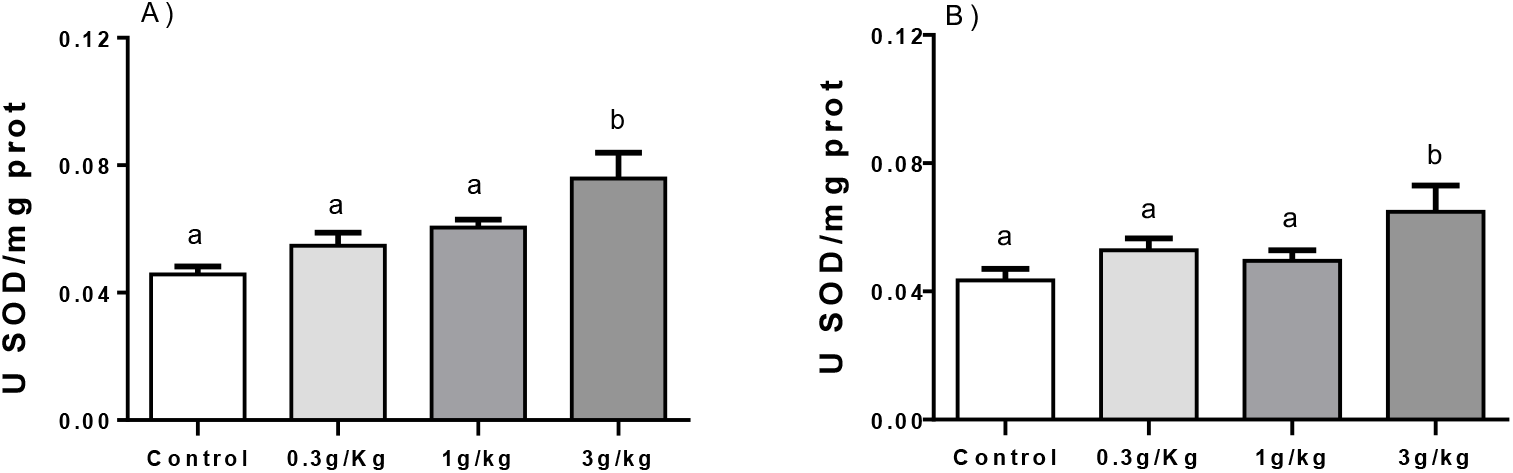
Effect of Caramel IV on SOD activity in liver (A) and kidney (B) of mice. Data are represented as U SOD/mg protein and expressed as mean ± S.E.M (n=9, p<0.05). Means for a variable with superscripts without a common letter differ.

#### 3.2.3 Acetylcholinesterase activity

All C-IV doses induced an increase in AChE activity when compared to the control of the liver (Fig. 7A, p<0.0001). On kidney, C-IV 3g/kg group increased AChE activity in comparison to group control (Fig. 7B, p<0.0055).

**Fig. 7.**
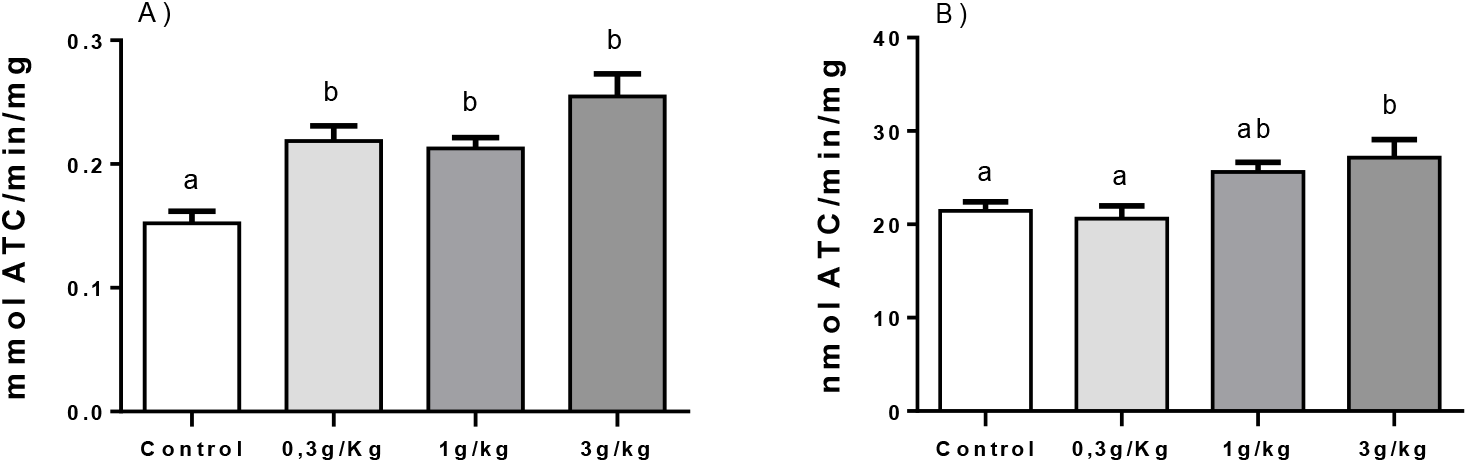
Effect of Caramel IV on AChE activity in liver (A) and kidney (B) of mice. Data are represented as nmol ATC/min/mg and expressed as mean ± S.E.M (n=9, p<0.05). Means for a variable with superscripts without a common letter differ.

### 3.3 Histological analysis

No histopathological changes were observed in control, 0.3g/kg and 1g/kg C-IV groups in both liver and kidney. However, figure 8D shows C-IV 3g/kg group induced a diffuse mild paracentral hydropic degeneration at the liver. Similarly, figure 8H demonstrate the presence of glomerulonephritis and moderate multifocal proliferative membrane and low glomerulus count caused by C-IV 3g/kg administration.

**Fig. 8.**
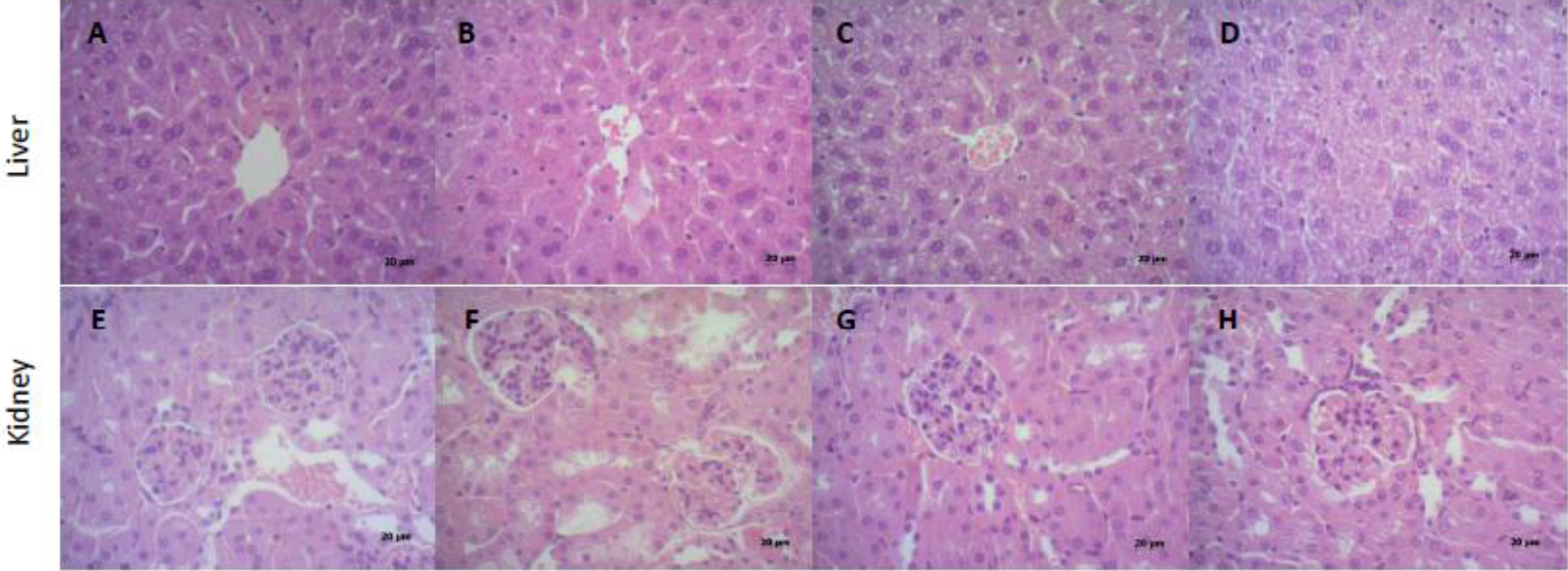
Photomicrographs (final magnification of 400x) in liver (A-D) and kidney (E-H) of mice by HE. Liver and kidney section of mice in the control, 0.3g/kg and 1g/kg C-IV showed normal structure of hepatocytes and glomerulus and renal tubules respectively. C-IV 3g/kg (D) shows a diffuse mild paracentral hydropic degeneration at liver. Kidney sections in C-IV 3g/kg group (H), present glomerulonephritis and moderate multifocal proliferative membrane and low glomerulus count.

## DISCUSSION

In this study, we demonstrated that C-IV 1g/kg and 3g/kg promoted the development of oxidative stress damages in mice tissues, represented by increased hepatic lipid peroxidation, reduction of NPSH groups and increased production of reactive species observed in renal tissue and histological alterations in both liver and kidney. Besides, we demonstrated that C-IV is responsible for activating the antioxidant defense system, represented by the increased activity of the enzymes catalase, superoxide dismutase and acetylcholinesterase on mice liver and kidney, but no effects on oxidative stress markers.

C-IV dye production process is characterized by the presence of nitrogen and sulfur in their manufacture chemical reactions, and its production leads to the high molecular weight constituents (HMW) formation (responsible for color differentiation) and low molecular weight constituents (LMW) formation (responsible for unintentional substances such as 4-methylimidazole (4-MeI)) ^5^. The effects of oxidative damage after C-IV administration here may be associated with 4-MEI, also classified as a human carcinogen and the most potent neurotoxic substance by the International Agency for Research on Cancer ^22^.

In this sense, mice treated with 4-MEI at 100, 130 and 160 g/kg presented a higher percentage of chromosomal mutations after 12 hours and 24 hours after 4-MEI exposure, indicating that 4-MEI is genocytotoxic to mice ^23^. Besides, studies evidenced increased neonatal mortality in rodents treated with C-IV, the formation of full moon neoplasms ^6^, tissue pigmentation alteration, and changes in body weight, in urine and hematological examinations ^10^.

The higher ROS production, especially hydrogen peroxide and superoxide, may be due to the mediation of cytochrome P450, which is recognized for promoting this increase ^24^, since 4-MEI has a structure very similar to 2 and 3-methylfuran which has the cytochrome P 450 protein after its metabolism ^25^. Similarly, high doses of paracetamol following cytochrome P450 metabolism generates a toxic metabolite known as N-acetyl-p-benzoquinoneimine, that can be conjugated with reduced glutathione and depleting its hepatic stores ^26^. We suggest that the cytochrome P450 pathway may be activated at high doses of C-IV due to the presence of 4-MEI, which is structurally similar to 2 and 3-methylfuran and has cytochrome P450 metabolism. Collaborating with this hypothesis, we demonstrated here that higher doses of C-IV are highly toxic through increased hepatic lipid peroxidation and may damage cell membranes, altering their structure and permeability ^27^.

In addition, glycotoxins, or lipid peroxidation products, promote the formation of advanced glycation and products (AGEs) ^28^ through the Maillard reaction, a process that occurs during the caramel dye caramelization process ^29^. C-IV is considered to be rich in AGEs, and their molecules are considered highly reactive acting as donors of electrons for the generation of free radicals, which can promote the development of oxidative stress and consequent cellular damage ^30^. AGEs activate specific receptors called RAGE (Receptor Advanced Glycation End products) and produce inflammatory cytokines, such as interleukin 1 and 6 (IL-1 and IL-6), tumor necrosis factor alpha (TNF-α), protein C reactive (PCR). This AGE-RAGE interaction seems to mediate the generation of free radicals, altering the permeability and vitality of the cellular membranes ^30^, which corroborates with our results about the formation of oxidative markers and leading to oxidative stress^28–30^.

The deleterious action of free radicals can be inhibited and/or reduced through the activation of the antioxidant defense system, maintaining the cellular homeostasis, preventing the amplification of the damages. According to our study, we demonstrated that C-IV was able to induce an increase in antioxidant enzyme defense system, represented by SOD, CAT, and acetylcholinesterase in the analyzed tissues ^31^. However, despite the antioxidant system modulation, it was not sufficient to prevent or blunt the oxidative damages induced by C-IV administration.

This oxidative stress damage could be observed by histological analysis in both liver and kidney of mice caused by C-IV 3g/kg administration, where diffuse mild paracentral hydropic degeneration could be identified at liver and a possible glomerulonephritis, moderate multifocal proliferative membrane, low glomerulus count at kidney tissue. In addition, studies show that animals treated with 2-methylimidazole develop renal lesions consisting of hemosiderin, which is most evident in male mice ^25^. These alterations are linked to increased hepatic lipid peroxidation and renal ROS damages.

This way, we point out that this is a critical study, since if we use a dosage conversion factor between human x mice – method that applies an exponent for body surface area, which account for difference in metabolic rate, to convert doses between animals and humans.- the highest dose use here is about to 30 fold lower than the RDA for humans ^32^. So, this highlight the relevance of our study, since low doses, even those below RDA, are causing many tissues damages and oxidative stress development.

Finally, it is known that AGEs production might lead to an inflammatory process ^30^, so, there is a high possibility that C-IV might lead to an inflammatory process, by increasing cytokines and triggering specific inflammation receptors ^30^. This hypothesis will be investigated in future studies from our group.

In conclusion, our study showed for the first time that high dose of Caramel Dye IV promotes effects on the oxidative profile, demonstrated by increased levels of TBARS in the liver, renal ROS production and NPSH reduction, given the potential cytotoxic effect of the same, since it possesses in its composition the substance 4-MEI, already being recognized for its harmful effects through possible routes of oxidation and metabolization. Besides, C-IV can promote a change in the redox system, observed by increased levels of renal CAT, SOD, and renal and hepatic acetylcholinesterase. Further studies are needed to understand the mechanisms that CI-V induces the oxidative and possible inflammation damages to cells.

## Acknowledgements

The work was supported by the PRONEM/CNPq/FAPERGS 16/25510000248-7 and CAPES/PROEX 23038.005848/2018-31 research grants. FAAS received a fellowship from CNPQ. JSS and PCR received a fellowship from CAPES.

## REFERENCES

(1) Prado, M. A.; Godoy, H. T. CORANTES ARTIFICIAIS EM ALIMENTOS. Alim. Nutr 2003. https://doi.org/10.1016/j.jpowsour.2008.03.048.

(2) Cunha, S. C.; Barrado, A. I.; Faria, M. A.; Fernandes, J. O. Assessment of 4-(5-)Methylimidazole in Soft Drinks and Dark Beer. J. Food Compos. Anal. 2011. https://doi.org/10.1016/j.jfca.2010.08.009.

(3) (ANS), E. P. on F. A. and N. S. added to F. Scientific Opinion on Safety of Steviol Glycosides for Proposed Uses as a Food Additive. EFSA J. 2010. https://doi.org/10.2903/j.efsa.2010.1537.

(4) Sengar, G.; Sharma, H. K. Food Caramels: A Review. J. Food Sci. Technol. 2012, 51 (9), 1686–1696. https://doi.org/10.1007/s13197-012-0633-z.

(5) Vollmuth, T. A. Caramel Color Safety – An Update. Food Chem. Toxicol. 2018, 111, 578–596. https://doi.org/10.1016/j.fct.2017.12.004.

(6) Program, N. T. Toxicology and Carcinogenesis Studies of 4-Methylimidazole (Cas No. 822-36-6) in F344/N Rats and B6C3F1 Mice (Feed Studies). Natl. Toxicol. Program Tech. Rep. Ser. 2007.

(7) Buschinelli, J. T. Manual Para Interpretação de Informações Sobre Substâncias Químicas; 2011.

(8) Seddon, M.; Looi, Y. H.; Shah, A. M. Oxidative Stress and Redox Signalling in Cardiac Hypertrophy and Heart Failure. Heart 2007, 93, 903–907. https://doi.org/10.1136/hrt.2005.068270.

(9) Stohs, S. J. The Role of Free Radicals in Toxicity and Disease. J. Basic Clin. Physiol. Pharmacol. 1995, 6 (3–4), 205–228. https://doi.org/10.1515/JBCPP.1995.6.3-4.205.

(10) MacKenzie, K. M.; Boysen, B. G.; Field, W. E.; Petsel, S. R. W.; Chappel, C. I.; Emerson, J. L.; Stanley, J. Toxicity and Carcinogenicity Studies of Caramel Colour IV in F344 Rats and B6C3F1mice. Food Chem. Toxicol. 1992, 30 (5), 431–443. https://doi.org/10.1016/0278-6915(92)90071-R.

(11) Ohkawa, H.; Ohishi, N.; Yagi, K. Assay for Lipid Peroxides in Animal Tissues by Thiobarbituric Acid Reaction. Anal. Biochem. 1979. https://doi.org/10.1016/0003-2697(79)90738-3.

(12) Myhre, O.; Andersen, J. M.; Aarnes, H.; Fonnum, F. Evaluation of the Probes 2′,7′-Dichlorofluorescin Diacetate, Luminol, and Lucigenin as Indicators of Reactive Species Formation. Biochemical Pharmacology. 2003. https://doi.org/10.1016/S0006-2952(03)00083-2.

(13) Pérez-Severiano, F.; Rodríguez-Pérez, M.; Pedraza-Chaverrí, J.; Maldonado, P. D.; Medina-Campos, O. N.; Ortíz-Plata, A.; Sánchez-García, A.; Villeda-Hernández, J.; Galván-Arzate, S.; Aguilera, P.; et al. S-Allylcysteine, a Garlic-Derived Antioxidant, Ameliorates Quinolinic Acid-Induced Neurotoxicity and Oxidative Damage in Rats. Neurochem. Int. 2004. https://doi.org/10.1016/j.neuint.2004.06.008.

(14) Ellman, G. L. Tissue Sulfhydryl Groups. Arch. Biochem. Biophys. 1959, 82 (1), 70–77. https://doi.org/10.1016/0003-9861(59)90090-6.

(15) Aebi, H. Catalase in Vitro. Methods Enzymol. 1984, 105, 121–126. https://doi.org/10.1016/s0076-6879(84)05016-3.

(16) Misra, H. P.; Fridovich, I. The Role of Superoxide Anion in the Autoxidation of Epinephrine and a Simple Assay for Superoxide Dismutase. J Biol Chem 1972, 247, 3170–3175.

(17) Bradford, M. M. A Rapid and Sensitive Method for the Quantitation of Microgram Quantities of Protein Utilizing the Principle of Protein-Dye Binding. Anal. Biochem. 1976. https://doi.org/10.1016/0003-2697(76)90527-3.

(18) Ellman, G. L.; Courtney, K. D.; Andres, V.; Feather-Stone, R. M. A New and Rapid Colorimetric Determination of Acetylcholinesterase Activity. Biochem. Pharmacol. 1961.

(19) Worek, F.; Mast, U.; Kiderlen, D.; Diepold, C.; Eyer, P. Improved Determination of Acetylcholinesterase Activity in Human Whole Blood. Clin. Chim. Acta 1999. https://doi.org/10.1016/S0009-8981(99)00144-8.

(20) Oliveira, V. A.; Favero, G.; Stacchiotti, A.; Giugno, L.; Buffoli, B.; de Oliveira, C. S.; Lavazza, A.; Albanese, M.; Rodella, L. F.; Pereira, M. E.; et al. Acute Mercury Exposition of Virgin, Pregnant, and Lactating Rats: Histopathological Kidney and Liver Evaluations. Environ. Toxicol. 2017, 32 (5), 1500–1512. https://doi.org/10.1002/tox.22370.

(21) Stacchiotti, A.; Favero, G.; Giugno, L.; Lavazza, A.; Reiter, R. J.; Rodella, L. F.; Rezzani, R. Mitochondrial and Metabolic Dysfunction in Renal Convoluted Tubules of Obese Mice: Protective Role of Melatonin. PLoS One 2014, 9 (10), e111141. https://doi.org/10.1371/journal.pone.0111141.

(22) Nishie, K.; Waiss, A. C.; Keyl, A. C. Toxicity of Methylimidazoles. Toxicol. Appl. Pharmacol. 1969. https://doi.org/10.1016/0041-008X(69)90111-2.

(23) Norizadeh Tazehkand, M.; Topaktas, M.; Yilmaz, M. B.; Hajipour, O.; Valipour, E. Delineating the Antigenotoxic and Anticytotoxic Potentials of 4-Methylimidazole against Ethyl Methanesulfonate Toxicity in Bone Marrow Cell of Swiss Albino Mice. Bratislava Med. J. 2016. https://doi.org/10.4149/BLL_2016_057.

(24) Du, K.; Ramachandran, A.; Jaeschke, H. Oxidative Stress during Acetaminophen Hepatotoxicity: Sources, Pathophysiological Role and Therapeutic Potential. Redox Biol. 2016, 10, 148–156. https://doi.org/10.1016/j.redox.2016.10.001.

(25) Chan, P. C.; Hills, G. D.; Kissling, G. E.; Nyska, A. Toxicity and Carcinogenicity Studies of 4-Methylimidazole in F344/N Rats and B6C3F1 Mice. Arch. Toxicol. 2008, 82 (1), 45–53. https://doi.org/10.1007/s00204-007-0222-5.

(26) Colle, D.; Arantes, L. P.; Gubert, P.; da Luz, S. C. A.; Athayde, M. L.; Teixeira Rocha, J. B.; Soares, F. A. A. Antioxidant Properties of Taraxacum Officinale Leaf Extract Are Involved in the Protective Effect Against Hepatoxicity Induced by Acetaminophen in Mice. J. Med. Food 2012, 15 (6), 549–556. https://doi.org/10.1089/jmf.2011.0282.

(27) Mello Filho, A. C.; Hoffmann, M. E.; Meneghini, R. Cell Killing and DNA Damage by Hydrogen Peroxide Are Mediated by Intracellular Iron. Biochem. J. 2015. https://doi.org/10.1042/bj2180273.

(28) Odani, H.; Shinzato, T.; Usami, J.; Matsumoto, Y.; Frye, E. B.; Baynes, J. W.; Maeda, K. Imidazolium Crosslinks Derived from Reaction of Lysine with Glyoxal and Methylglyoxal Are Increased in Serum Proteins of Uremic Patients: Evidence for Increased Oxidative Stress in Uremia. FEBS Lett. 1998, 427 (3), 381–385. https://doi.org/10.1016/S0014-5793(98)00416-5.

(29) Nseir, W.; Nassar, F.; Assy, N. Soft Drinks Consumption and Nonalcoholic Fatty Liver Disease. World J. Gastroenterol. 2010, 16 (21), 2579–2588. https://doi.org/10.3748/wjg.v16.i21.2579.

(30) Voyer, L. E. ARTÍCULO ESPECIAL REACCIÓN DE MAILLARD. EFECTOS PATOGÉNICOS Glicación Endógena Efectos Patogénicos. 2019, 137–143.

(31) Barbosa, K. B. F.; Costa, N. M. B.; Alfenas, R. de C. G.; De Paula, S. O.; Minim, V. P. R.; Bressan, J. Estresse Oxidativo: Conceito, Implicações e Fatores Modulatórios. Rev. Nutr. 2010, 23 (4), 629–643. https://doi.org/10.1590/S1415-52732010000400013.

(32) Nair, A. B.; Jacob, S. A Simple Practice Guide for Dose Conversion between Animals and Human. J. basic Clin. Pharm. 2016. https://doi.org/10.4103/0976-0105.177703.

